## Supplementary figures and images for "Cerebrospinal fluid-contacting neuron tracing reveals structural and functional connectivity for locomotion in the mouse spinal cord"

### Figure S

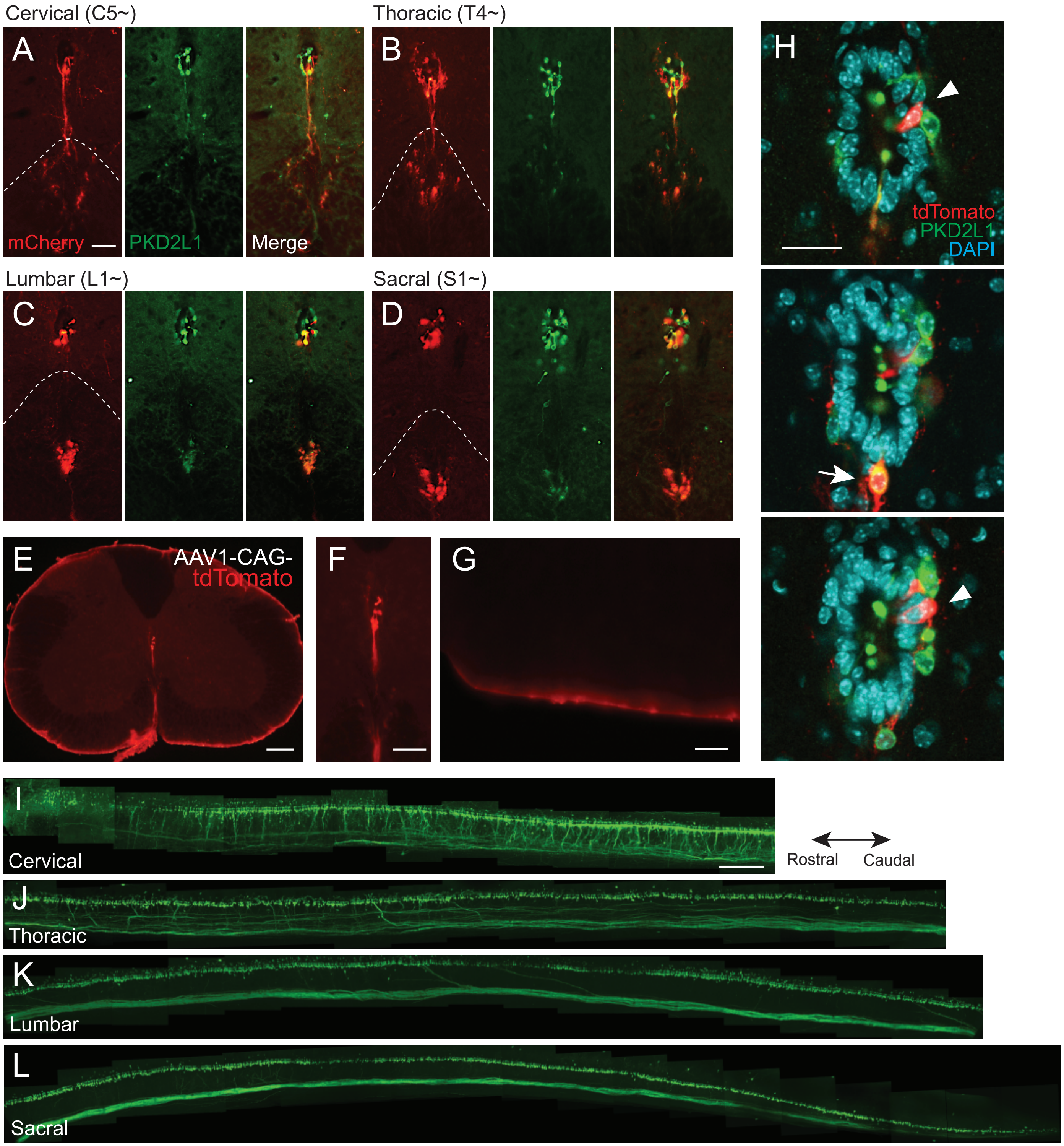

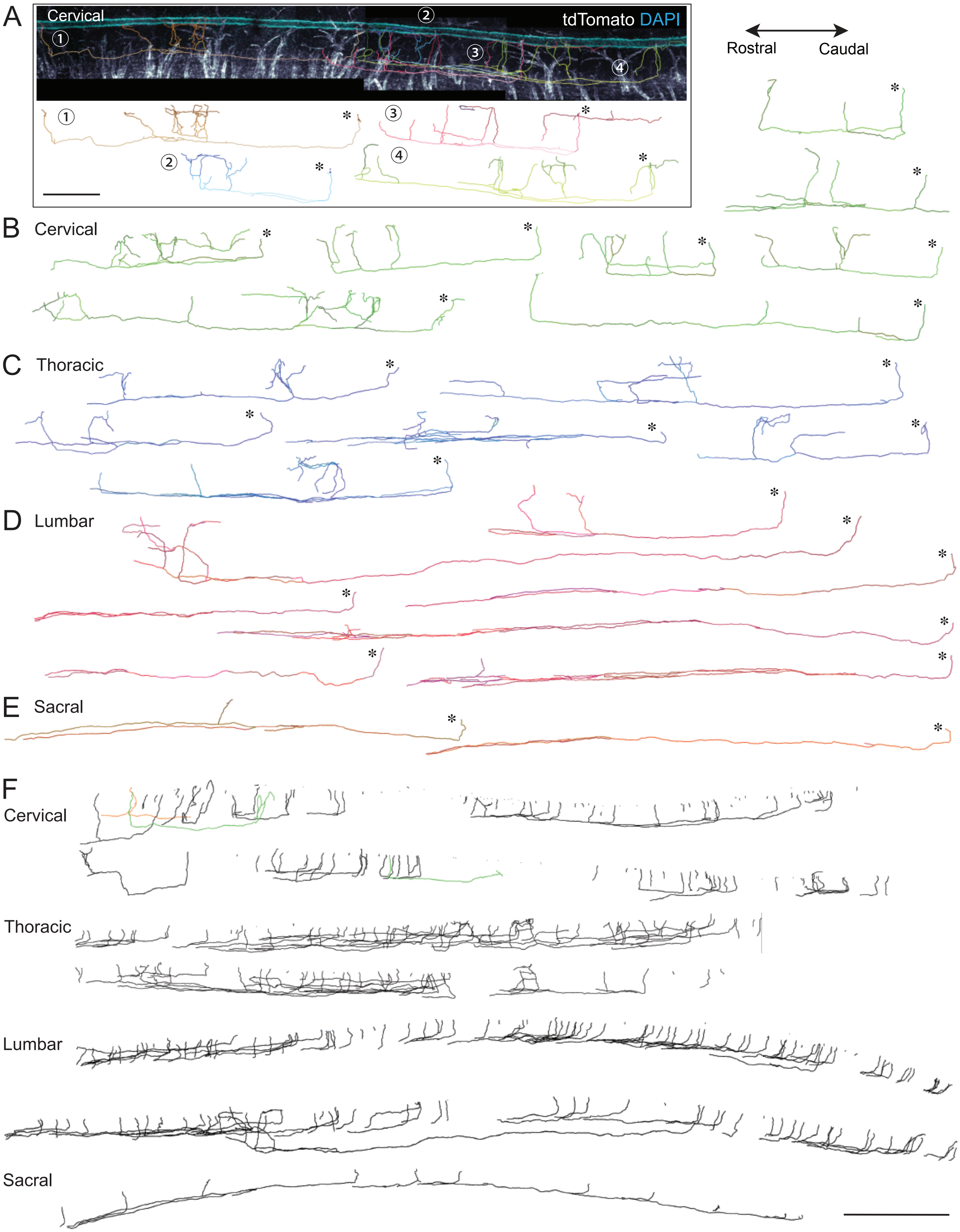

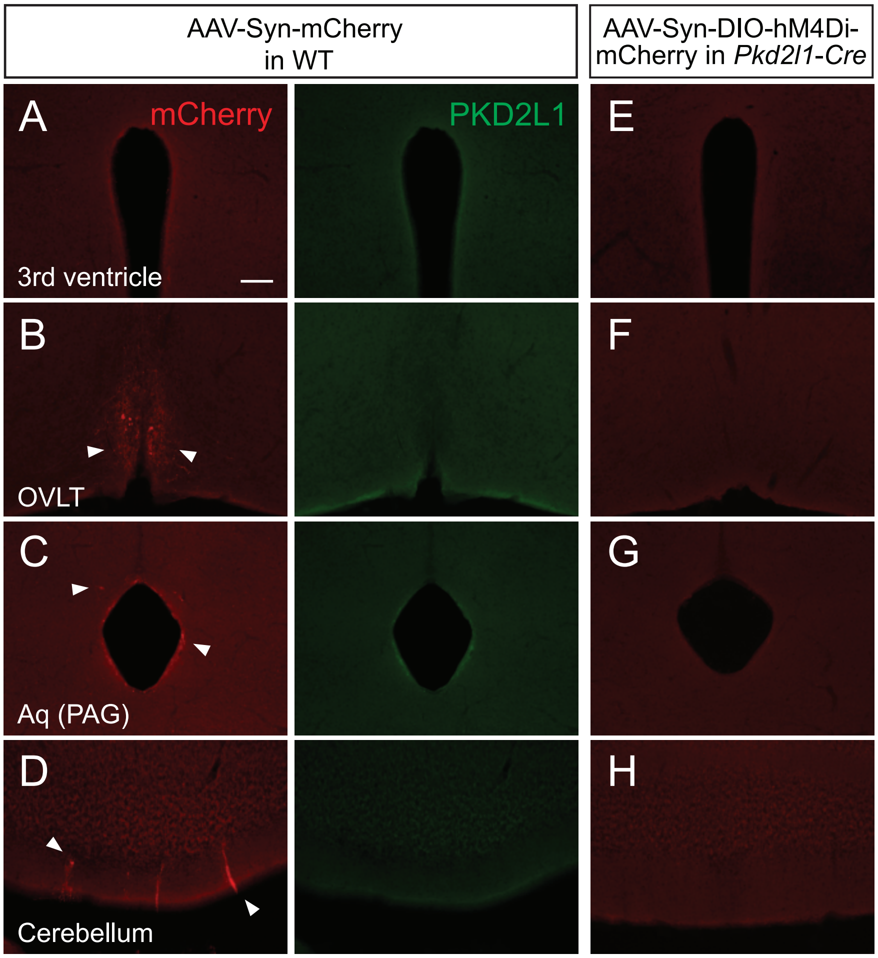

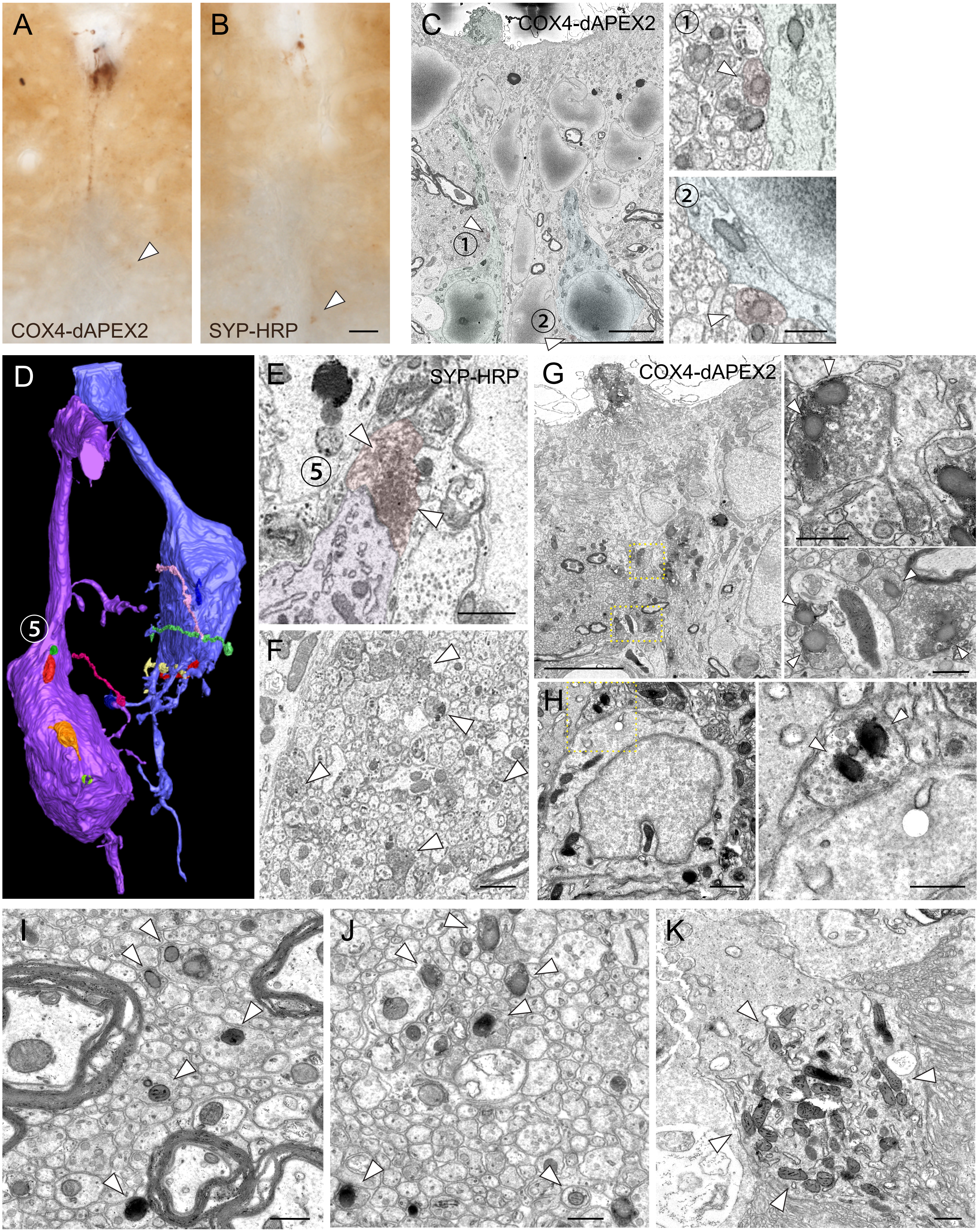

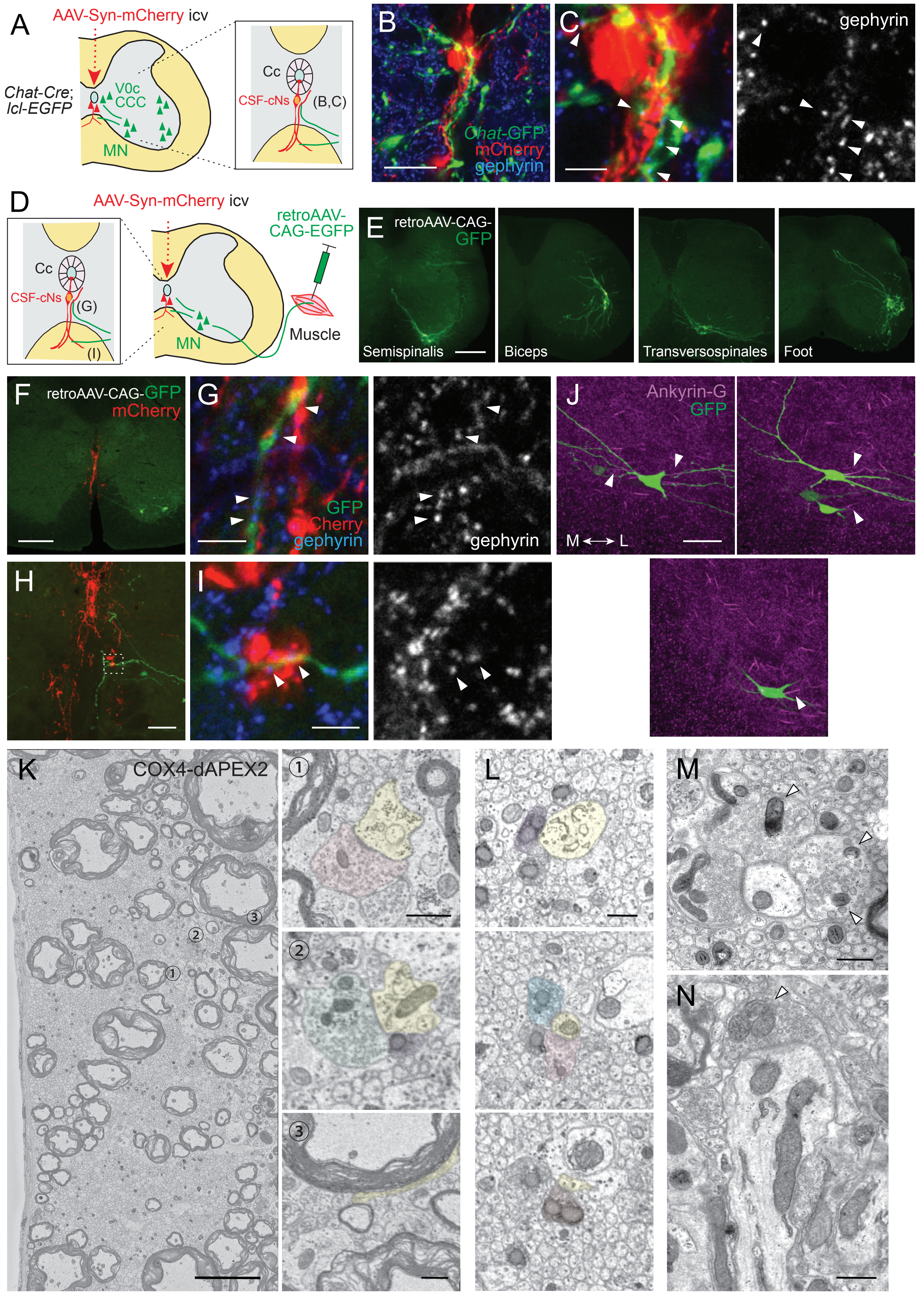

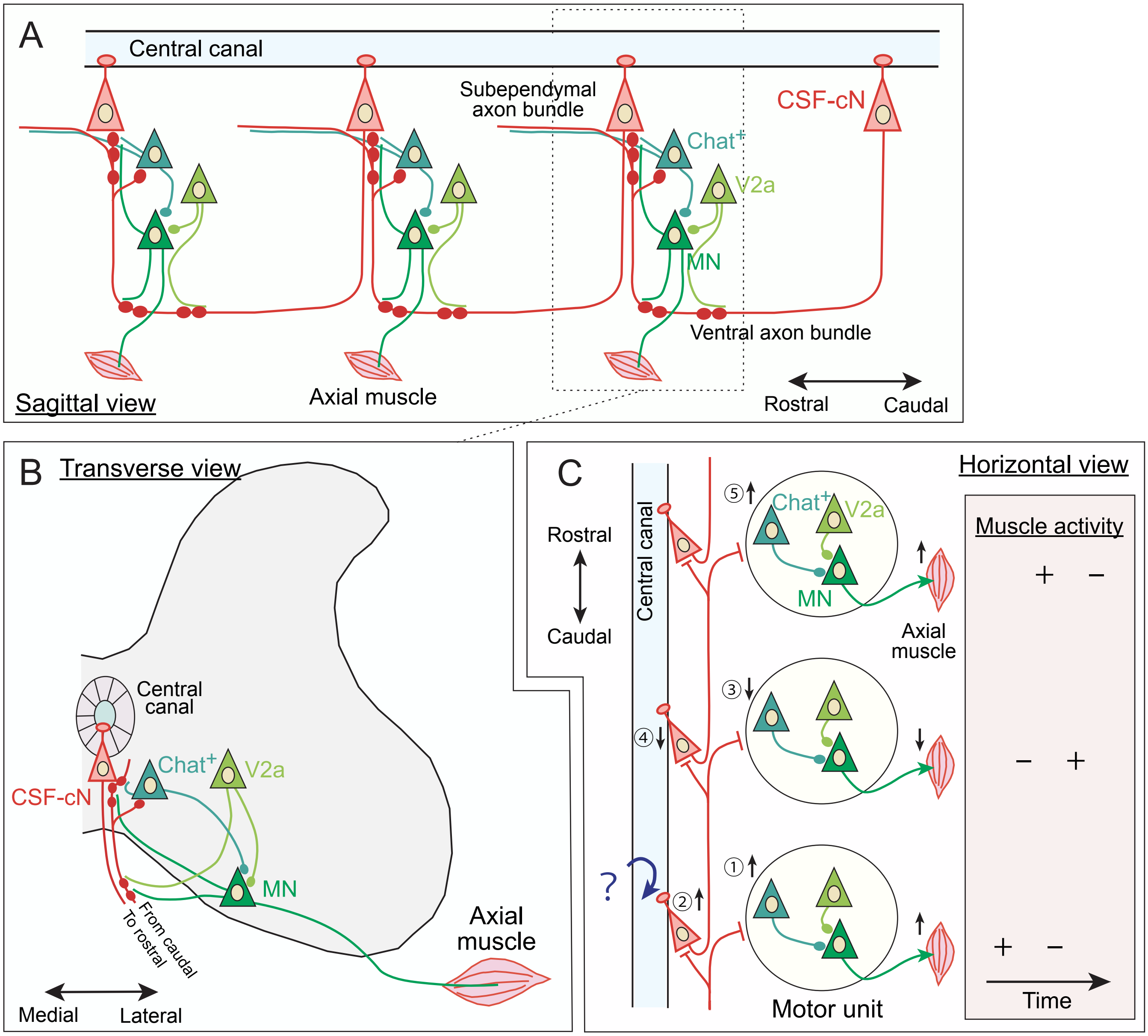
